# Going beyond cell clustering and feature aggregation: Is there single cell level information in single-cell ATAC-seq data?

**DOI:** 10.1101/2024.12.04.626927

**Authors:** Aaron Wing Cheung Kwok, Heejung Shim, Davis J McCarthy

**Affiliations:** Bioinformatics and Cellular Genomics, St Vincent’s Institute of Medical Research, Fitzroy, VIC 3065, Australia; Melbourne Integrative Genomics, University of Melbourne, Parkville, VIC, 3010, Australia; School of Mathematics and Statistics, Faculty of Science, University of Melbourne, Parkville, VIC, 3010, Australia; Faculty of Medicine, Dentistry and Health Sciences, University of Melbourne, Parkville, VIC, 3010, Australia

## Abstract

Single-cell Assay for Transposase Accessible Chromatin with sequencing (scATAC-seq) has become a widely used method for investigating chromatin accessibility at single-cell resolution. However, the resulting data is highly sparse with most data entries being zeros. As such, currently available computational methods for scATAC-seq feature a range of transformation procedures to extract meaningful information from the sparse data. Most notably, these transformations can be categorized into: 1) feature aggregation with known biological associations, 2) pseudo-bulking cells of similar biology, and 3) binarisation of count data. These strategies beg the question of whether or not scATAC-seq data actually has usable single-cell and single-region information as intended from the assay. If we can go beyond aggregated features and pooled cells, it opens up the possibility of more complex statistical tasks that require that degree of granularity. To reach the finest possible resolution of single-cell, single-region information there are inevitably many computational challenges to overcome. Here, we review the major data analysis challenges lying between raw data readout and biological discovery, and discuss the limitations of current data analysis approaches. Lastly, we conclude that chromatin accessibility profiling at true single-cell resolution is not yet achieved with current technology, but that it may be achieved with promising developments in optimising the efficiency of scATAC-seq assays.

## 2 Introduction

Single-cell Assay for Transposase Accessible Chromatin with sequencing (scATAC-seq) has established itself as one of the most popular assays for interrogating chromatin accessibility at single-cell resolution [1]. The assay relies on Tn5 tranposase which simultaneously fragments accessible DNA regions and integrates adapter sequences, during a process termed ‘tagmentation’ [2]. The DNA fragments from each single cell are then sequenced and quantified which serves as the entry point for data analysis. However, computational analyses of said data is exceptionally challenging due to the data readout of scATAC-seq being sparse, with over 90% of the entries in the count matrix being zeros [3]. This challenge motivates the development of a plethora of novel computational tools to answer meaningful questions about chromatin accessibility. Here, we describe a typical computational workflow for analysing scATAC-seq data and the major challenges associated with each step (Fig. 1). Starting from the initial readout i.e.

**Figure 1:**
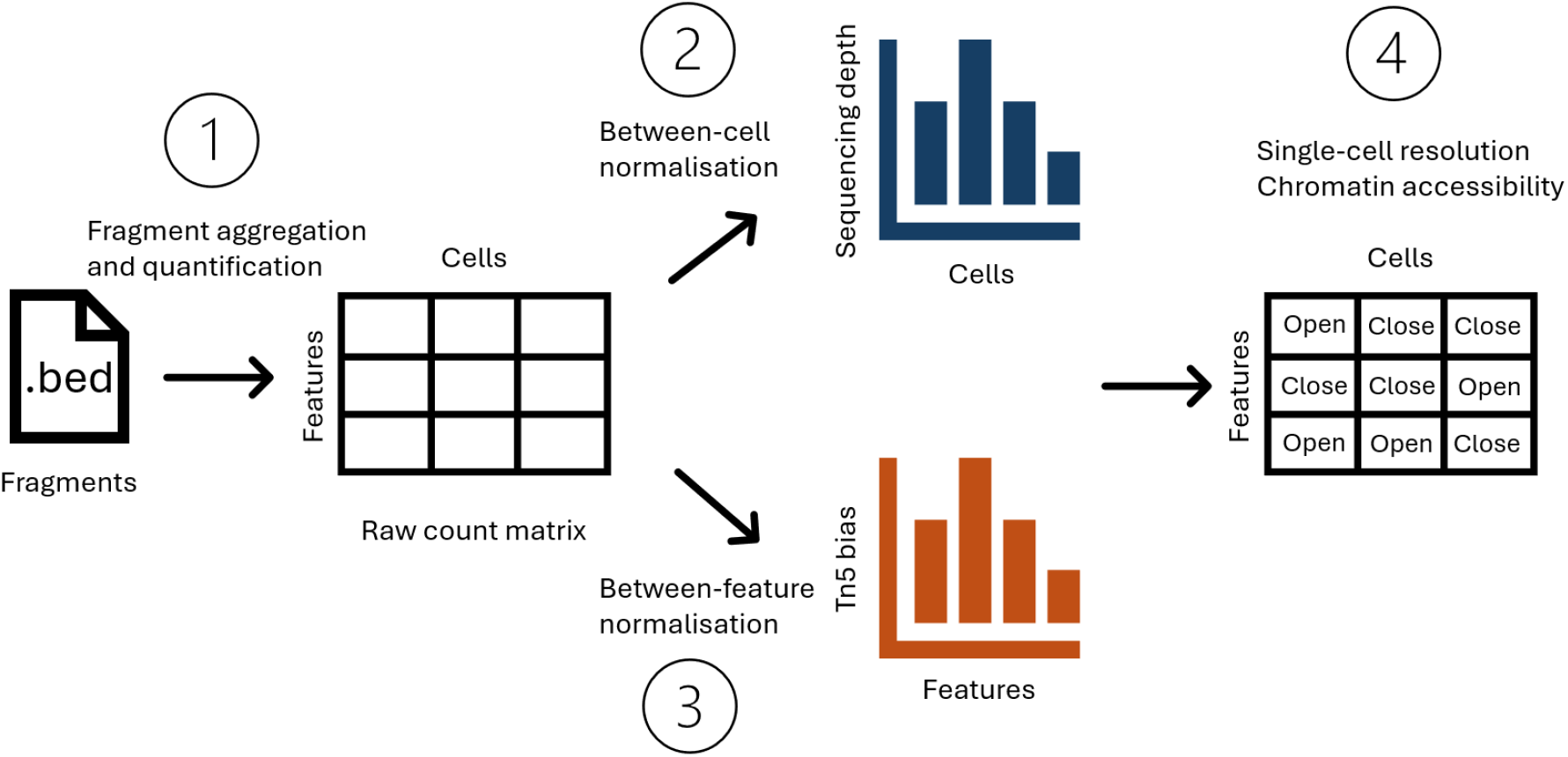
Conceptual diagram for key challenges in typical scATAC-seq data analysis, including 1) fragment aggregation and quantification; 2) Between-cell normalisation; 3) Between-feature normalisation; and 4) Interpreting chromatin accessibility at single-cell resolution.

DNA fragments, feature engineering is necessary to group fragments from the whole genome into regions of interest. Using this set of regions of interest, a count matrix can be obtained for various downstream analysis tasks. Next, normalisation is typically performed to remove between-cell and/or between-region technical biases, which is usually followed by dimension reduction. Using low-dimensional representations, more concrete biological questions can be addressed, such as cell type annotation, differential accessibility, and motif enrichment.

Although these computational steps are highly analogous to single-cell transcriptome analyses, extreme data sparsity presents unique challenges at each stage of analysis. Below, we elaborate on 4 major challenges in this typical pipeline that remain largely unsolved, with little consensus on the best way to approach them.

### 2.1 Major challenge 1: Fragment aggregation and quantification

Like other single-cell modalities, most analysis workflows for scATAC-seq data start with a count matrix. However, quantifying chromatin accessibility is not as straightforward as quantifying gene expression. First, genomic features for scATAC-seq are ambiguous and not standardised, unlike in transcriptomics where features are defined by well-annotated genes and transcripts. In scATAC-seq analyses, researchers will either divide the whole genome into fixed-width windows or identify signal-enriched regions using peak callers to limit the analysis to biologically relevant regions of interest. The choice is usually up to users’ preferences, but occasionally determined by the strategy employed by a specific computational tool. Secondly, within the defined features (be it fixed-width windows or called peaks), whether to count individual Tn5 insertion events or the presence of whole fragments is another topic up for debate. As a result, for the same raw fragment file, different counting strategies can generate different count matrices.

These intricacies are discussed in great detail by Miao and Kim [4], who propose paired insertion counts (PIC) as the preferred quantification method for scATAC-seq data. The advantage of using PIC is two-fold: it has attractive statistical properties for modeling purposes; and as pointed out by Miao and Kim [4] and Martens et al. [5], the quantitative nature of scATAC-seq readout can be related to biology. As such, here we opt to frame our discussion around PIC quntification of chromatin accessibility from scATAC-seq data. For simplicity, we will use fixed-width (500bp) regions to avoid having to account for variable peak sizes.

### 2.2 Major challenge 2: Sequencing depth normalisation

Sequencing depth variation between cells is a common source of unwanted variation in any single-cell sequencing data. If not properly accounted for, the variation in sequencing depth can be the largest source of between-cell variation and mask biological heterogeneity. In single-cell RNA-seq (scRNA-seq) data analyses, variation in sequencing depth is usually dealt with via normalisation prior to downstream analysis. For scATAC-seq data, the most widely used option is Term Frequency-Inverse Document Frequency (TF-IDF) normalisation. It is implemented with different flavours in popular tools such as Signac [6], ArchR [7], scOpen [3], and Cell Ranger ATAC [8] (summarised in Table 1). Importantly, TF-IDF preserves the region *×* cell dimensions of the count matrix without prior aggregation. However, benchmark studies show that it is often ineffective in removing library size effects [9] and, despite its popularity, there is little discussion on why that is the case. As such, choosing a particular TF-IDF flavour is mostly based on heuristics, personal preferences, and default settings in software packages.

**Table 1:** Comparison of different flavours of TF-IDF implementation.

| Method | TF-IDF | Features | Counting | Binarise |
| --- | --- | --- | --- | --- |
| Signac [6] | $\log(\text{TF} \times 10^4 \times \text{IDF} + 1)$ | Peaks | Fragments | No |
| ArchR [7], scOpen [3] | $\text{TF} \times \log(\text{IDF} + 1)$ | 500bp bins | Insertions | Yes |
| Cusanovich [10] | $\text{TF} \times \log(\text{IDF} + 1)$ | 5kb bins | Insertions | Yes |
| Hill [11] | $\log(\text{TF} \times 10^5 + 1) \times \log(\text{IDF})$ | Peaks/bins | Insertions | Yes |
| Cell Ranger ATAC [8] | $\log(\text{IDF})$ | Peaks | Insertions | No |

### 2.3 Major challenge 3: Region-specific bias

Detection of open chromatin with ATAC-seq heavily relies on the tagmentation activity of Tn5 enzyme, which has a preference for some genomic sequence characteristics over others, leading to technical variation between regions that does not necessarily reflect differences between local accessibility [12]. For bulk ATAC-seq, strategies have been developed to mitigate the effect of Tn5 cleavage bias on downstream analysis, such as weight matrix scaling (ATACorrect [13]), position dependency models (HINT-ATAC [14]), and k-mer based methods like SELMA [15]. Apart from sequence composition, it has been shown that epigenetic features such as DNA motif, shape, and methylation can drive Tn5 preferences [16]. The overall mechanism of Tn5 bias is complex and difficult to quantify accurately. Therefore, to reduce the scope of this study, we chose to showcase GC-content as a representative for region-specific bias, which is a well known factor that drives sample-specific technical bias in DNA sequencing (DNA-seq), Chromatin Immunoprecipitation sequencing (ChIP-seq), and RNA sequencing (RNA-seq) data [17]. For bulk ATAC-seq, normalisation with regard to GC-content is also crucial to avoid confounding downstream analysis [18]. Although the same effects should be expected in scATAC-seq as well, there is rarely a bespoke step in pipelines to normalise for GC effects, unless when some aggregation has been done beforehand that amplifies technical bias, e.g., chromVAR aggregating peaks based on motifs [19].

### 2.4 Major challenge 4: Interpretation of chromatin accessibility

Despite being the main motivation for scATAC-seq, the interpretation of ‘profiling chromatin accessibility at single-cell resolution’ is unclear. A longstanding notion treats chromatin accessibility as binary: a region is either open or closed in a cell. In reality, with two copies of each chromosome in a cell (for autosomes in a diploid organism), the ‘true’ chromatin accessibility state is at least ternary: both chromosomes open, both closed, or one closed and one open. Moreover, recent studies showed that scATAC-seq counts have quantitative information instead [4, 5], as biological factors such as nucleosome turnover rate can contribute to the quantitative observation of chromatin accessibility [5]. To take it even further, it has recently been argued that it is unclear whether euchromatin should even be considered ‘open’ *per se* [20]. As such, depending on the biological assumptions, the interpretation of ‘chromatin accessibility at single-cell resolution’ can vary and thus introduces ambiguity when interrogating scATAC-seq data. These nuances are rarely addressed as most computational analyses are limited to cluster or cell type level, where counts are aggregated and treated as if they were continuous, like gene expression.

To truly realise the ‘single-cell’ in scATAC-seq, the major challenges reviewed above must be addressed. Here, we first show that common ad-hoc normalisation methods are ineffective in removing technical biases from scATAC-seq data, due to both theoretical and practical reasons. To deal with technical biases while avoiding arbitrary transformations, we propose a hierarchical count model to infer single-cell level chromatin information. Chromatin biology is highly complex, therefore to reduce the scope of the study, we aim first to establish the simplest case, which is to assume chromatin accessibility is binary at a single cell-single region level, i.e., a cell is either open or closed for a single region. Using the proposed model, we find that current scATAC-seq count data in general does not have sufficient information to perform such inference. Thus, we can currently only rely on aggregation as a temporary solution for scATAC-seq data analyses. However, assays that optimize for Tn5 sensitivity show promising results and represent the best path toward achieving true single-cell resolution.

## 3 Results

### 3.1 TF-IDF approaches are counterproductive in removing sequencing depth biases

To explain the poor benchmark performances from TF-IDF based methods [21], we will elaborate on its calculation and theoretical limitations. As the name suggests, TF-IDF is the product of two distinct parts: Term Frequency (TF) and Inverse Document Frequency (IDF). Here, we unpack the two parts of TF-IDF and identify inherent limitations in its application as a default normalisation strategy for sequencing depth variation in scATAC-seq data.

#### 3.1.1 Term Frequency

We work with an N *×* P ‘count matrix’ X which holds information about the number of observed counts in N cells and P features. The features can represent either peaks or bins depending on the upstream data pre-processing approach. We let i ∈ {1, …, N} index cells and j ∈ {1, …, P} index features, so that x_ij_ is the observed count of the jth feature in the ith cell.

The term frequency transformation of a particular count value is defined as the count value divided by the sum of counts over all features in the same cell as the count value,

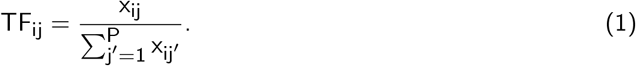

We can compare this value to counts per ten thousand (CPTT) commonly used in scaling scRNA-seq counts:

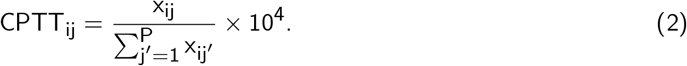

Clearly, these two quantities are identical except for the scaling factor of 10^4^. In RNA-seq terminology, it is equivalent to counts per million (CPM) divided by 100. The smaller scaling factor here is used to account for the overall smaller library sizes observed in single cell assays compared to bulk.

Dividing by total count is a sound strategy for bulk sequencing as the read counts are often in the magnitudes of hundreds to thousands, with total counts per sample in the millions. However, in scATAC-seq data, most data entries share the same value at either 0 or 1 (comprising of 90-95% of the data), but the total count of each cell is different. Therefore, after TF transformation, the largest variation between cells will naturally be due to their denominators, that is, the total counts per cell or sequencing depth (Fig. 2a). This effect is further exacerbated by binarising the counts before transformation (as done in some popular analyses software, e.g., ArchR, scOpen), which forces all non-zero entries to share the same value of 1 (Fig. 2a). Ironically, the aim of this strategy is to remove sequencing depth variation, but it ends up introducing extra information about library sizes instead.

**Figure 2:**
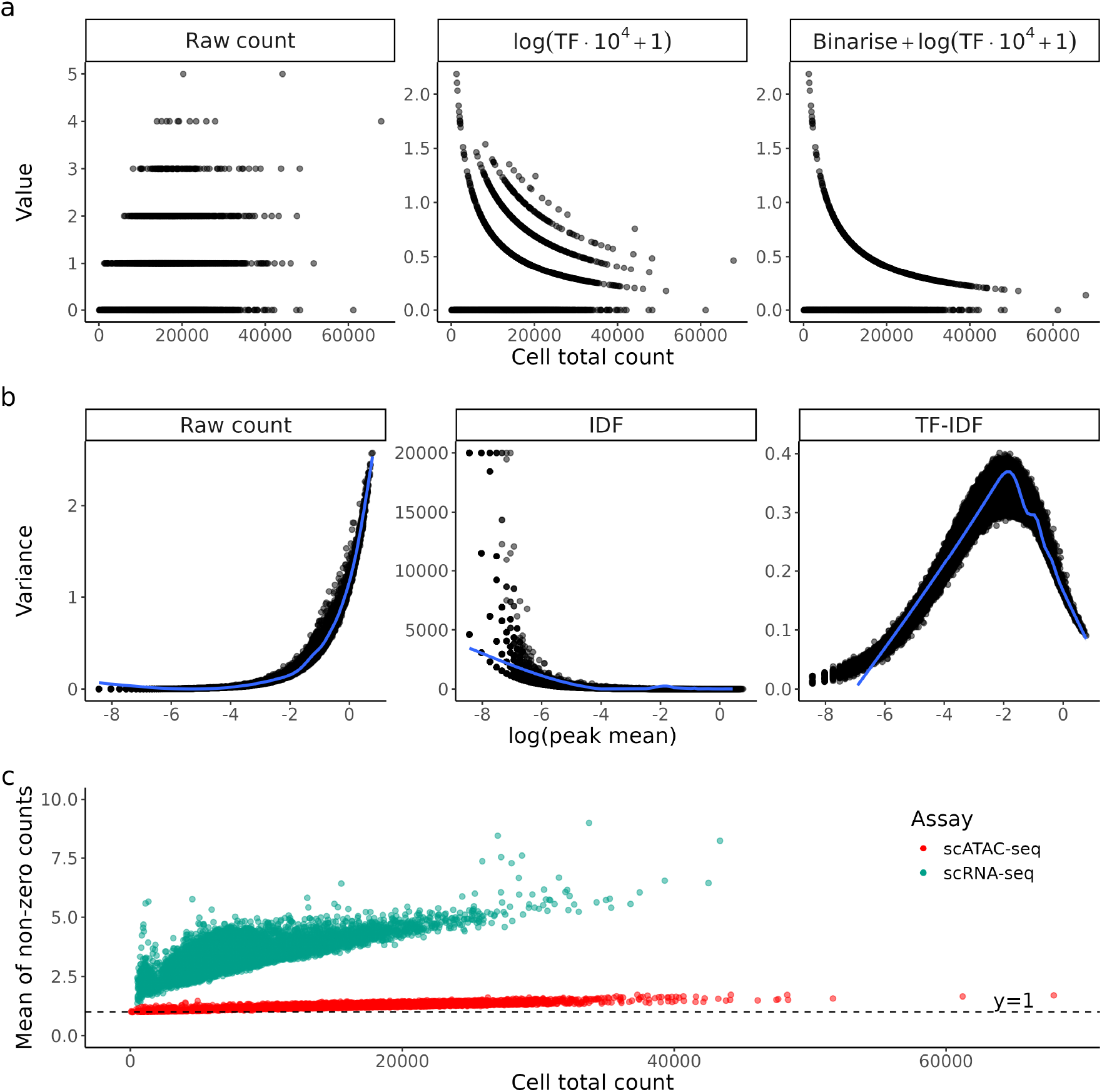
a) Raw counts and their TF-transformed values for a random region in PBMC10k scATAC-seq dataset, plotted against the total count of each cell. Each dot is a cell. Here the region chr1:1273633-1274133 was chosen as demonstration. b) Variance of raw count and IDF-transformed values plotted against mean of raw count of each region. Each dot is a region. c) Mean of non-zero counts in each cell plotted against its total count for both scRNA-seq and scATAC-seq PBMC10k.

Due to the large number of genomic regions and likely small number of Tn5 cuts in each region, the majority of observed counts of scATAC-seq data is exactly zero (Fig. S1). Thus, an increasing sequencing depth will more likely turn a 0 into 1 instead of turning a 1 to a value larger than 1. We observed that the mean of non-zero counts in scATAC-seq rarely go above 1.2 even in cells with high total counts, which is on average 62.8% lower than that of scRNA-seq data (Fig. 2c). In other words, sequencing depth difference is mostly represented by sparsity and normalisation methods that target non-zero values (e.g., dividing by total count/a linear size factor) will not address the problem effectively. This has been a known issue for scRNA-seq, where bulk-based methods like log(CPM+ 1) were found to be sub-optimal as they fail to account for exact zeros and the arbitrary choice of pseudocount can introduce subtle bias to the data [22]. TF transformation, being a rehash of log(CPM + 1), suffers from the same issues as its scRNA-seq counterpart as we observe parallels in count characteristics.

#### 3.1.2 Inverse Document Frequency

IDF is a feature-wise metric that weights features according to their rarity among all features, given by:

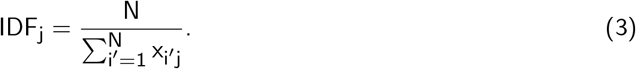

We can also rewrite IDF in terms of region mean count *µ*_j_:

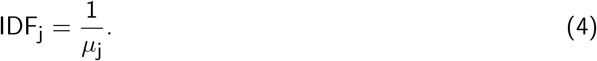

The intuition behind IDF is to give more weight to regions that are rarely open as they are more likely to correlate with cell type specific functions, while less weight is given to regions that are open in most cells as they are likely to be involved in housekeeping functions that are not relevant to cell type. In a normal cell clustering task, this weighting scheme is sensible, but it should not be viewed as a typical ‘normalisation’ technique that can transfer to other tasks. Dividing all count values from a region with a region-specific constant *µ*_j_ introduces additional dependency between variance and mean (Fig. 2b). To be specific, the variance will be scaled by a factor of 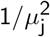. Caution has to be exercised when applying IDF transformed counts to models that assume uniform variance as IDF will inherently tend to exacerbate heteroskedasticity in scATAC-seq data.

### 3.2 GC correction methods designed for bulk ATAC-seq do not transfer well to single cell

The effect of GC-content on bulk ATAC-seq readout is well characterized [18] and we observed the same effect on scATAC-seq data (Fig. 3a), where regions with higher GC-content tend to have higher mean counts, with the effect varying between cell types. While such a relationship can be explained by biology due to many accessible regions being gene promoters which often have high GC-content [23], technical variation between regions will make them hard to compare and possibly confound analyses that involve region-to-region comparison. Unfortunately, as speculated by Van den Berge et al. [18], GC-aware normalisation methods for bulk ATAC-seq have limited utility on its single-cell counterpart. We tried 2 recommended methods: GC-full-quantile normalisation (GC-FQ) and smooth GC-FQ (Section 6.2.2) on a subset of cell types from the hematopoietic cells dataset [8]. We found that quantile-based methods that performed well on bulk ATAC-seq data do not have a significant effect on the overall relationship between GC-content and mean count (Fig. 3b,c), although the disparity between cell types is reduced for GC-rich peaks. For GC-FQ, it even comes at the cost of increased sparsity as the median peak mean drops by an order of magnitude in general (Fig. 3b).

**Figure 3:**
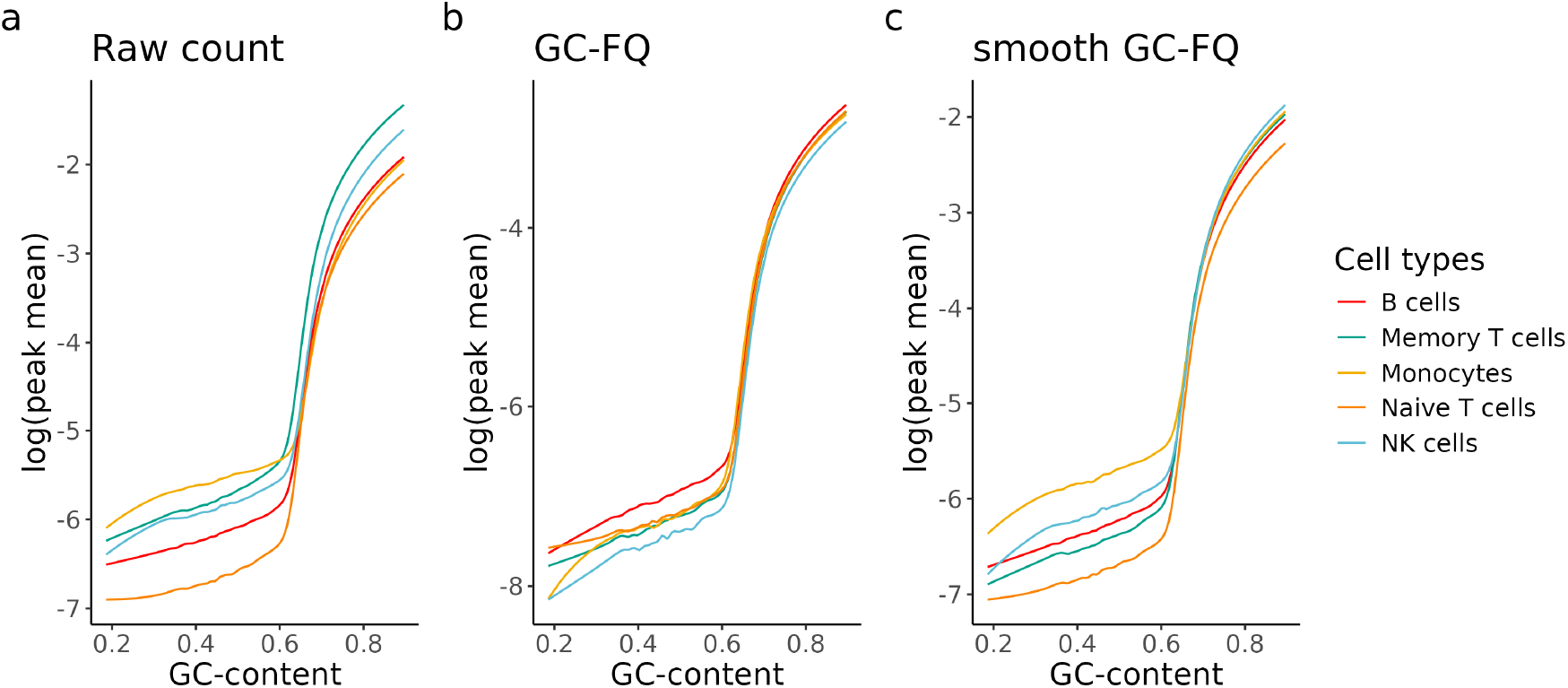
Fitted lowess curves of log(mean count) as a function of GC-content for 5 of the annotated cell types in the hematopoietic cells dataset [8]. The fit was performed on a) raw counts, b) GC-FQ normalised counts, and c) smooth GC-FQ normalised counts. Note that the shape and slope of the curves are different for different cell types.

### 3.3 A hierarchical model for inferring cell chromatin states

With the intention of dealing with all the technical biases listed above and also inferring per-cell, perregion open/closed information from scATAC-seq data, we constructed the following hierarchical model. We work with a N *×* P scATAC-seq paired insertion count (PIC) matrix [4]. Let i = {1, …, N} index cells and j = {1, …, P} index chromatin regions. We define a mixture model that describes the observed count x_ij_ with the following hierarchical structure:

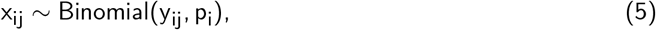

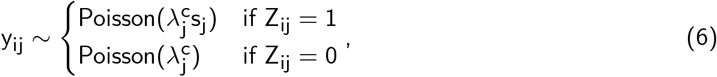

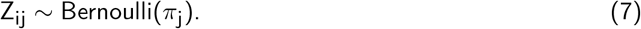

Where:

p_i_ denotes cell-specific observation probability;

y_ij_ denotes true number of paired Tn5 cuts (latent);

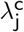 denotes count rate for closed cells (background count rate due to GC effect);

s_j_ denotes signal-to-noise ratio;

*π*_j_ denotes proportion of open cells for a given region.

The motivation for this model specification is to describe biological and technical processes with explainable variables. We have aimed to keep the model as simple, and thus as interpretable, as possible while capturing the most important aspects of the data generation process. We stick to the notion that for a given region, single cells can either be open or closed, as discussed in Section 2.4. The proportion of open cells is denoted by *π*_j_. In an scATAC-seq experiment, DNA regions are fragmented depending on their accessibility state (Z_ij_), affinity for Tn5 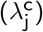, and signal-to-noise ratio (s_j_), but not every accessible region in every cell can be fragmented by Tn5. This property is represented by the Poisson distribution. Lastly, the resulting latent fragments are subjected to technical sampling bias that varies among cells, which is represented by the binomial distribution.

Our model addresses the previously stated major challenges as follows:

- Modelling counts instead of binarised data to extract more information, as suggested by Miao and Kim [4] and Martens et al. [5]. This approach is not inherently contradictory to the assumption of chromatin accessibility being a binary trait. Intuitively, a higher fragment count should indicate a higher confidence of the cell being ‘open’ in a region and vice versa.
- Our modelling approach has the advantage of retaining the region *×* cell dimension of the count matrix and requires no arbitrary transformation or prior clustering and cell type annotation.
- Inferring binary state of each cell (open/closed) through using the posterior probability of Z_ij_, i.e., P(Z_ij_ = 1|data).
- Instead of using total count as a scaling factor, using the binomial observation probability p_i_ is a more faithful representation of fragment dropout. This approach is conceptually similar to the observation probability in the PIC model (Methods 6.3.2) [4].
- Specifying a background rate 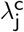 to be region-specific accounts for region-specific biases such as

GC-content variation. In theory, one can further specify 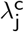 to be a function of any known technical effect. In our analysis we chose GC-content to be the representative region-specific effect.

We will apply this simple model to address the key challenges in scATAC-seq data outlined above and draw conclusions about current approaches to modelling and analysing scATAC-seq data.

### 3.4 Current scATAC-seq data does not have enough information to infer single-cell level chromatin state

The lack of ground truth makes it difficult to properly evaluate our model on real datasets. Therefore, we first simulated data with a wide range of parameters to: 1) quantify the level of information needed to perform accurate inference, and 2) get a rough idea of how real data would behave.

We simulated 10,000 cells from our hierarchical model with varying background rates 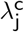 and signal-to-noise ratios s_j_ (Methods 6.3). We estimated p_i_ from data (Methods 6.3.2) and fixed *π*_j_ to 0.3 for demonstration purposes. Our findings are mostly invariant to the choice of *π*_j_ (Fig. S2). For each simulated scenario, P(Z_ij_ = 1 | x_ij_) is calculated using ground truth parameters (Methods 6.3.6). The posterior is then evaluated against the ground truth chromatin state of each observation. Each scenario was repeated 30 times and evaluated by the mean Area Under Receiver Operating Characteristic Curve (AUROC) (Fig. 4a). Given that the counts were simulated from the same model and ground truth parameters were used to compute the posterior, we would expect the posterior to be highly informative for identifying cells that are ‘open’, i.e., high AUROC. The opposite case would suggest that there is a ‘component collapse’ problem, i.e., open cells and closed cells do not have a significant difference in counts and cannot be told apart.

**Figure 4:**
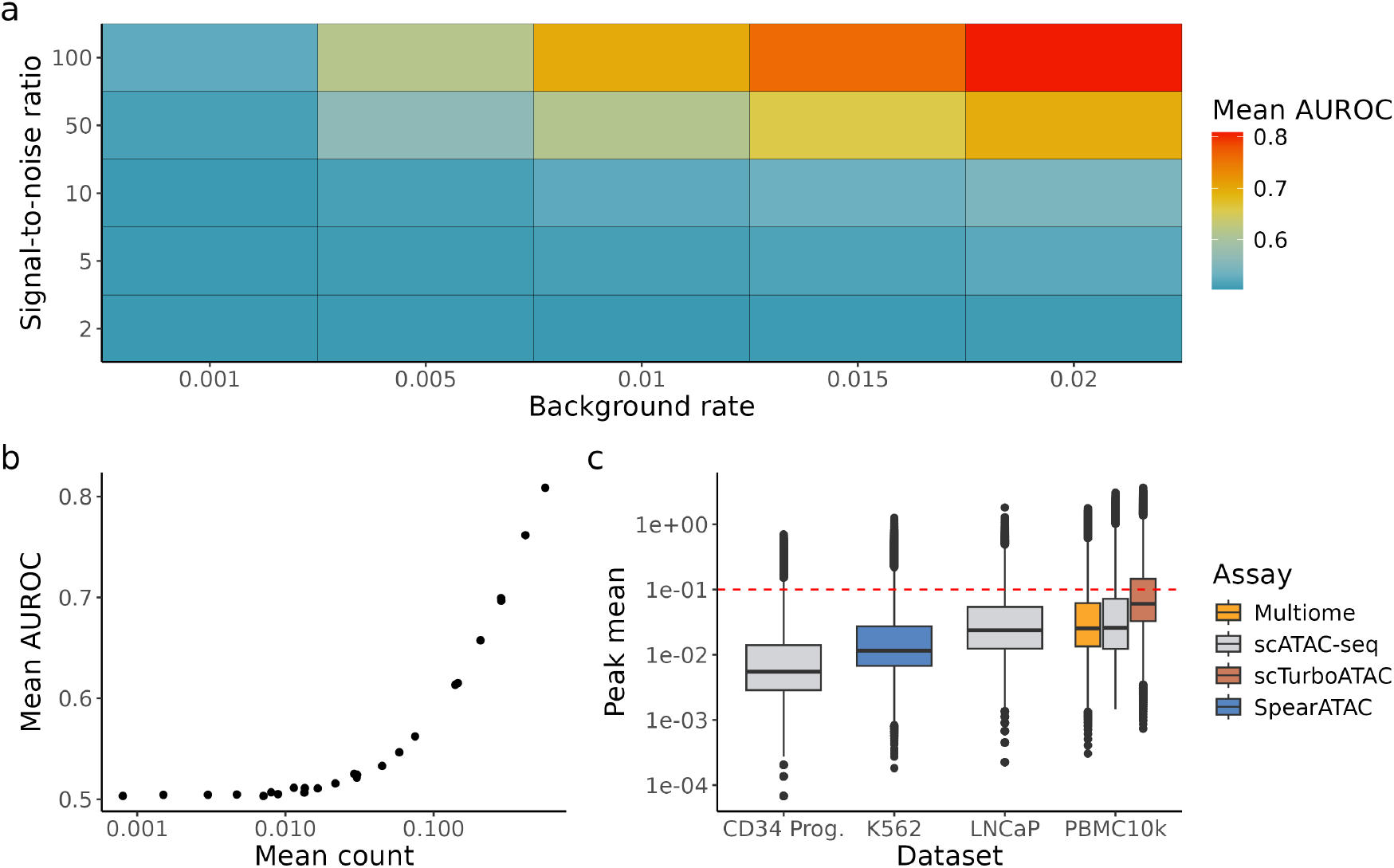
a) Simulation data with different combinations of background rates and signal-to-noise ratio. *π*_j_ is fixed to 0.3 for demonstration purposes. For each scenario, simulation is repeated for 30 times and the mean AUROC is calculated. b) Mean AUROC against mean of simulated counts. c) Box plot of peak mean from 6 datasets with varying biology and assays. Red dotted line marks the point where mean count *≥* 0.1, corresponding to AUROC *≥* 0.55 in our simulations.

Even with perfect retrieval of parameters, chromatin states are almost unidentifiable in situations with low 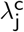 or low s_j_ (Fig. 4a), indicating a severe lack of information in these simulated scenarios. The best case scenario is when both parameters are high 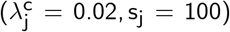, with mean AUROC 0.84. We also found that classification performance correlates strongly with mean count (Fig. 4b), i.e., it is in general easier to correctly identify chromatin states of single cells in peaks with higher counts, which is intuitive as mean is a function of 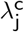 and s_j_. This result can serve as a practical guidance as one cannot directly observe the underlying parameters in real data. When comparing the mean count of real data against that of simulated counts, we found that in most datasets, less than 25% of peaks have sufficient counts to resemble simulation scenarios with mean AUROC 0.55 or higher (Fig. 4c), indicating more than 75% of features likely have insufficient information to infer chromatin states in single cells. However, scTurboATAC [24], an scATAC-seq assay optimised for Tn5 sensitivity, generated more fragments than other datasets, with 34% of peaks having mean count higher than 0.1 which corresponds to mean AUROC *≥* 0.55 in our simulations (Fig. 4c).

Although the majority of peaks have low information, biologists are often most interested in a small subset of biologically significant peaks (e.g., peaks that are strongly associated with marker genes) as they are highly informative in cell type identification. Therefore we also asked how much single-cell level information these marker peaks hold, by extending our simulation analysis to better reflect real biology. To incorporate prior biological knowledge to our simulations, we estimated parameters from the hematopoietic cells scATAC-seq dataset from Satpathy et al. [8]. Briefly, we first estimated the cell-specific observation probability p_i_ using the PIC model (Methods 6.3.2). Then, we estimated 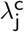 based on the GC-content of closed chromatin regions (Methods 6.2) and specified *π*_j_ to be the cell type proportion based on published annotations (Methods 6.3.4). Finally, s_j_ can be solved by matching the first moment (Methods 6.3.5). Counts were then simulated using these parameters and the posterior was evaluated in the same way as the synthetic data analysis (Methods 6.3.6).

We chose 9 peaks that are close to 9 well-known marker genes (Methods 6.3.5, Table 4) as their accessibility is expected to be highly correlated with the abundance of their respective cell type, e.g., a peak that is close to the *CD19* gene body should be accessible in most B cells. Simulation based on real data shows that most marker peaks have sufficient information at the single-cell level with performance positively correlated to mean count (Fig. 5a), which is consistent with our fully synthetic simulation. Notably, even for a prominent marker peak like *CD19* with a 94th percentile mean count across all peaks, its signal is too low to infer chromatin state at a single-cell level (mean AUROC *≈* 0.55).

**Figure 5:**
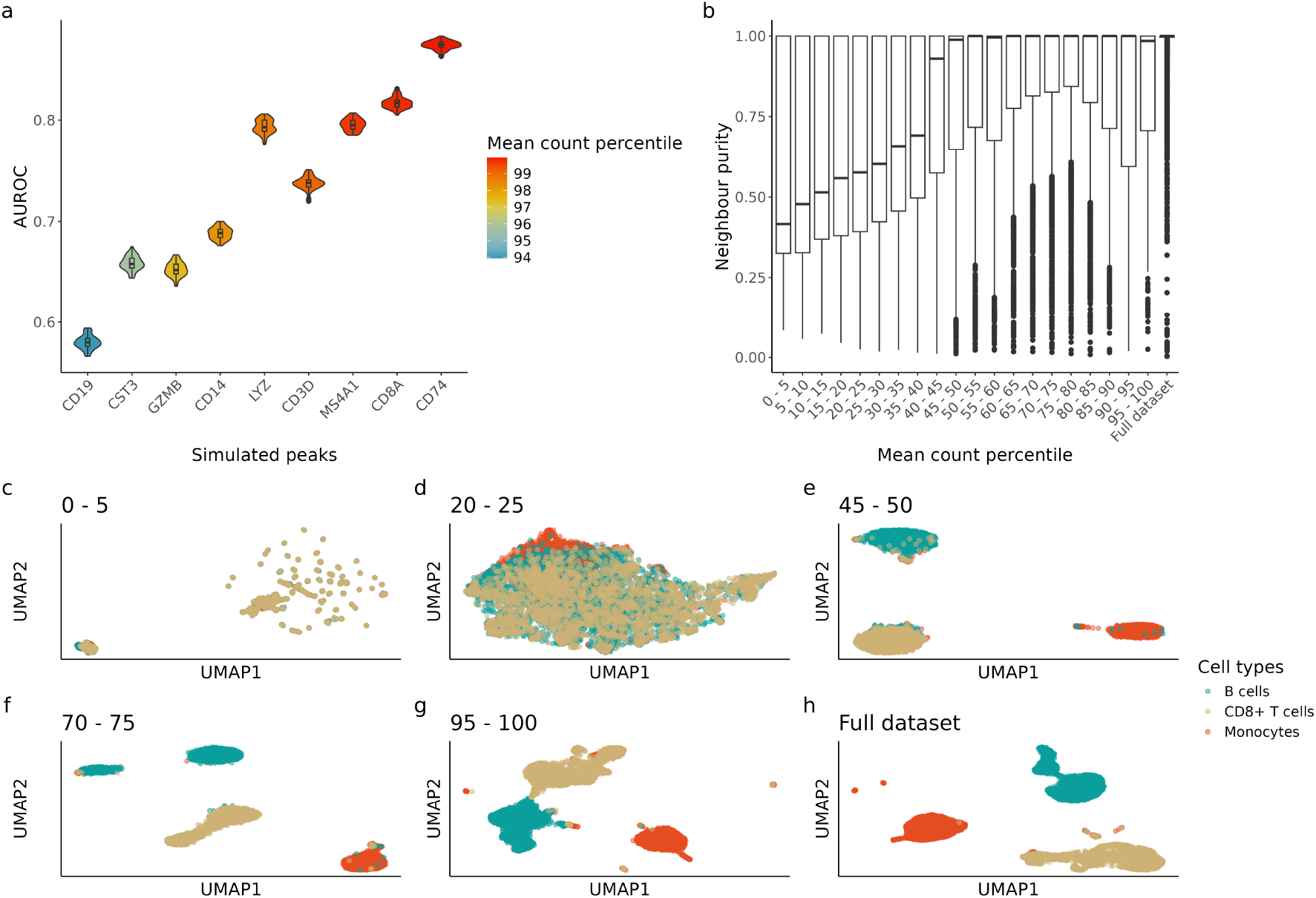
Simulation based on hematopoietic cells dataset. a) AUROC of simulated peaks based on marker genes. Simulated peaks are ordered based on mean count percentile of the whole peak set (571,400 peaks) where parameters were estimated from. b) Neighbour purity calculated using top 30 LSI components calculated with different peak quantiles (Section 6.4). c-h) Uniform manifold approximation and projection (UMAP) visualisation of cell embeddings calculated using different peak quantiles (Section 6.4). Only a selection of UMAPs are shown here, for clustering visualisation using each peak quantile please refer to Fig. S3.

### 3.5 Aggregation and dimensionality reduction can serve as temporary solutions

Another way to aggregate information across features is through dimension reduction, which is pivotal to most scATAC-seq analysis pipelines. Principal component analysis (PCA) uses information from all features to produce orthogonal axes that explain the most variance, effectively aggregating features in a softer sense than directly summing up counts. It is often recommended to filter for the top 5-10% peaks before performing dimension reduction as opposed to first finding highly variable genes (HVGs) in scRNA-seq. We hypothesised that this heuristic works in general because peaks with sufficient signal for single cell chromatin state inference (i.e., peaks with top 5-10% mean count) are the major contributors to the low dimensional space. Thereby, without the high count peaks, the dimension-reduced data should retain a lot less information and thus greatly affect subsequent tasks that depend on the representation, such as clustering.

To assess the contribution of peaks with different level of counts on clustering performance, we subset peaks into 20 quantiles according to their mean counts and obtained their low dimension embeddings using one quantile of peaks at a time (Methods 6.4). Briefly, to get the embeddings, each quantile of peaks is subject to Latent Semantic Indexing (LSI), i.e., TF-IDF followed by Singular Value Decomposition (SVD). Despite its pitfalls (previously discussed in Section 3.1), LSI is the most popular dimension reduction approach for scATAC-seq data, so it is relevant to show how feature selection impacts a typical clustering pipeline. Then, we evaluated the quality of the embeddings by calculating the neighbour purity of each cell with the ‘bluster’ R package [25]. Briefly, for each cell, its 50 nearest neighbours are identified using the embeddings and the fraction of neighbouring cells sharing the same cell type label is measured. Higher neighbourhood purity indicates higher degree of separation between cell types, which indicates a better low dimension representation. Under our hypothesis, embeddings obtained from the top peak quantiles should have significantly higher neighbour purity than their lower count counterparts.

As expected, clustering with the top 5% peaks (95th-100th percentile) gives very similar results to clustering with all peaks (Fig. 5b,g,h). However, clustering with any quantile above the median also gives similar results to clustering with all peaks, with the median neighbour purity being close to 1 and improvement between quantiles starting to diminish (Fig. 5b). There are two insights from this result: 1) although most peaks have insufficient information on a single-cell level, biology can be effectively recovered from the aggregate via dimension reduction; 2) clustering with the top 5-10% peaks is hardly better than some other sets of peaks with lower mean count, suggesting that the typical feature selection procedures can risk losing information and including a larger subset of peaks for cell type clustering can be beneficial. This corroborates with the benchmark findings from Luo et al. [9] as they found both ArchR and SnapATAC2 benefit from using a larger subset of features than default.

## 4 Discussion

We presented a hierarchical count model that is motivated by the data generating process of scATAC-seq data. However, we showed with various simulations that current scATAC-seq data is too sparse to infer true single cell chromatin states. While this result might be due to limitations in our assumptions about chromatin accessibility, we reason that if scATAC-seq does not have enough information to recover the simplest binary case, then it is highly likely that more complicated biological models (e.g., ternary, quantitative chromatin states) are also unrecoverable. As such, it appears that to claim we have succeeded in profiling chromatin accessibility at single-cell resolution would be a misnomer.

However, meaningful biology can still be extracted from scATAC-seq data on a cell type level with appropriate analysis approaches that account for technical biases. To that end, we recommend against treating TF-IDF as a depth normalisation method due to theoretical limitations shown. While we do not deny its utility in tasks such as cell type clustering, the resulting counts are not ‘depth-normalised’. In many cases, the sequencing depth effect is even exaggerated after TF-IDF transformation, leading to yet another bandage solution in analyses, specifically, removing the first principal component manually before clustering. Single-cell transcriptomic data analyses have multiple available methods for depth normalisation. In contrast, apart from TF-IDF, there is a lack of methods for scATAC-seq data analyses that simply return ‘depth-normalised counts’ with the same dimensions. However, there tools not based on TF-IDF that incorporate sequencing depth information into downstream tasks without explicitly normalising with total count or size factor. Instead, they try to learn the relationship between sequencing depth and observed count directly from the data. For example, PeakVI [26] trains a neural net specifically on learning the cell-specific scaling factor; and PACS [27] parameterises the sequencing depth effect as an observation probability which is learnt from count data directly. A recent benchmark [21] also showed that linear regression-based normalisation implemented in SnapATAC [28] is more robust for more difficult clustering tasks.

As a workaround for the lack of resolution in scATAC-seq, aggregation is necessary to extract useful information. Current common practice in scATAC-seq data analysis is to aggregate information from similar features for downstream tasks. One way to do it is by summing up counts from similar peaks, with the similarity often defined by genomic features. For example, chromVAR [19] groups peaks by the presence of certain motifs, effectively reducing the sparse peak *×* cell matrix into a smaller motif *×* cell matrix. Whereas, BROCKMAN [29] summarises peaks based on k-mer frequencies around the insertion sites and Cicero [30] summarises peaks at the gene level by calculating gene activity scores. We showed that dimension reduction as a softer form of aggregation is effective in cell clustering, but the common practice of selecting features with top 5-10% mean count does not show significant improvement in clustering over using some feature subsets with lower count. Although these low count peaks have minimal information individually, they might still be valuable in a reduced dimension space.

Apart from aggregating features, another approach we have not analysed is to increase signal by aggregate biologically similar cells prior to analysis. The traditional way is to pseudo-bulk cell types and aggregate by either sum or mean, but the concept of ‘metacells’[31] as a finer grain version of cell type clusters should also be considered. However, the concept of open or closed is ambiguous as the resolution is no longer single cell and a cell aggregate can contain an arbitrary number of open cells. In this case, a model that treats chromatin accessibility as a quantitative trait, such as the PACS model [27], might be more suitable. Another concern cell-type or metacell aggregation would be its dependence on low dimension embeddings. Many cell clustering algorithms, including meta-cell methods like SEAcell [32], rely on the constructing a k-nearest neighbour (KNN) graph from low dimension embeddings, which in turn relies on proper data preprocessing and normalisation. How best to preprocess and normalise are still open questions for scATAC-seq data where the most recommended LSI method has statistical pitfalls and the prevalent assumption of binary-ness is challenged [4, 5]. Similar to feature aggregation, perhaps a softer form of aggregation instead of hard assignment to groups can be considered to boost signals in individual cells.

No matter how sophisticated computational methods get, ultimately the chromatin accessibility information that can be extracted from scATAC-seq is largely limited by the efficiency of Tn5 transposase insertion [24]. Our previous simulations show that it is possible to reliably infer cell chromatin states within a peak given a sufficient amount of information. However, this can only be achieved by improving the sensitivity of Tn5 transposase itself such that more insertion events can happen. One such example is scTurboATAC [24], in which the sensitivity and versatility of Tn5 transposase were enhanced with optimised experimental workflows. Though this does not guarantee a significant increase in single-cell level information, we believe an experimental approach to address the enormous sparsity in scATAC-seq data is a step to the right direction. Future assay improvements should strive to not only increase signal but also minimise noise to optimise for a better signal-to-noise ratio.

## 5 Conclusion

To conclude, we have provided a general overview of problems in scATAC-seq data analysis, such as fragment quantification, normalisation, and interpretation of of ‘chromatin accessibility’. In particular, we show that the widely used TF-IDF normalisation has statistical pitfalls that exacerbate technical bias. We proposed a hierarchical model to infer single-cell chromatin states from scATAC-seq counts. However, our simulation shows that with the sparsity in current scATAC-seq data, it is almost impossible to accurately identify whether a cell is open or closed in a chromatin region. Although this lack of resolution can be circumvented with aggregation and dimension reduction to obtain meaningful biological results from scATAC-seq data, measurement of chromatin accessibility at true single-cell resolution is still far from being achieved. To realise this goal, improving the sensitivity of scATAC-seq assays appears to be a promising avenue.

## 6 Methods

### 6.1 Datasets and preprocessing

#### 6.1.1 Downloading data

All datasets used in this study are publicly available (Table 2). The PBMC10k datasets (scATAC-seq, scRNA-seq, and Multiome) were downloaded from the 10X Genomics website. (Link to scATAC-seq, scRNA-seq, Multiome). The fragment files for the hematopoietic cells dataset [8] were downloaded from GEO with accession number GSE129785. Processed data object with cell barcodes, called peak set, and cell type annotations (scATAC Heme All SummarizedExperiment.final.rds) was downloaded from github (https://github.com/GreenleafLab/10x-scATAC-2019). The fragment files and processed data objects with cell barcodes, called peak set, and cell type annotations for K562 SpearATAC dataset [33] were downloaded from GEO with accession number GSE168851. The fragment files for LNCaP dataset [34] were downloaded from GEO with accession number GSE168667. The fragment files for PBMC10k scTurboATAC dataset [24] were downloaded from GEO with accession number GSE235506.

**Table 2:** Summary of datasets used.

| Dataset | Assay | Citation | Accession | # cells | # features |
| --- | --- | --- | --- | --- | --- |
| PBMC10k | scATAC-seq | 10X Genomics | 10X website | 10,246 | 191,833 |
| PBMC10k | scRNA-seq | 10X Genomics | 10X website | 11,922 | 22,302 |
| PBMC10k | Multiome | 10X Genomics | 10X website | 9,829 | 160,216 |
| PBMC10k | scTurboATAC | Seufert et al. [24] | GSE235506 | 8,128 | 243,114 |
| Hematopoietic cells | scATAC-seq | Satpathy et al. [8] | GSE129785 | 63,882 | 571,400 |
| LNCaP | scATAC-seq | Taavitsainen et al. [34] | GSE168667 | 4,436 | 112,049 |
| K562 | SpearATAC | Pierce et al. [33] | GSE168851 | 32,832 | 277,112 |

#### 6.1.2 Peak calling and generating PIC matrices for scATAC-seq data

We used the R package ‘PICsnATAC’ v(1.0.0) [4] to generate PIC matrices. The PIC counting() function requires 3 inputs: 1) fragment file, 2) cell barcodes, and 3) peak set. For datasets with uniform-length peak sets available (hematopoietic cells dataset and K562 SpearATAC dataset), the called peak set and cell barcodes were directly used as input along with the downloaded fragment files. For datasets with non-uniform-length peak sets or no peak set available, we obtained cell barcodes and peak set by running the default ArchR (v1.0.3)[7] pipeline with the downloaded fragment files as input. For the reference genome, we followed the version that was used to produce the fragment files (Table 3). We filtered cells using default parameters for (minTSS = 4; minFrags = 1000). We then called 500bp peaks using the addReproduciblePeakSet() function with MACS2 as the backend. The resulting cell barcodes and peak set were used as input to generate PIC matrices.

**Table 3:** Reference genome version used for each scATAC-seq dataset.

| Dataset | Genome |
| --- | --- |
| PBMC10k scATAC-seq | hg38 |
| PBMC10k Multiome | hg38 |
| Hematopoietic cells | hg19 |
| PBMC10k scTurboATAC | hg38 |
| LNCaP | hg38 |
| K562 | hg38 |

**Table 4:** Marker genes used.

| Cell type | Marker genes | Citation | Chosen peak |
| --- | --- | --- | --- |
| CD8+ T cells | CD8A | Uhlen et al. [36] | chr2:87013156-87013656 |
|  | GZMB |  | chr14:25142649-25143149 |
|  | CD3D |  | chr11:118209254-118209754 |
| B cells | CD19 | Uhlen et al. [36], Karlsson et al. [37] | chr16:28941954-28942454 |
|  | MS4A1 |  | chr11:60223007-60223507 |
|  | CD74 |  | chr5:149790045-149790545 |
| Monocytes | CD14 | Uhlen et al. [36] | chr5:140012001-140012501 |
|  | LYZ |  | chr12:69723632-69724132 |
|  | CST3 |  | chr20:23636332-23636832 |

### 6.2 GC-content normalisation

#### 6.2.1 GC-content retrieval

We used the Bioconductor R package Biostrings (v2.70.3) [35] to retrieve the GC-content of every peak region, using the reference genome of the relevant dataset. Table 3 provides the genome version used for each dataset.

#### 6.2.2 Normalisation methods

We adapted code from Van den Berge et al. [18] to test bulk ATAC-seq normalisation methods on scATAC-seq data. We tested 2 GC-aware methods that performed well in their benchmark: GC-full-quantile normalisation (GC-FQ) and smooth GC-FQ normalisation. Briefly, they are both based on full-quantile normalisation, which features 1) sorting the counts for each cell, 2) replacing all elements of each feature with its median, then 3) unsorting each cell. For more details on these methods, please see Van den Berge et al. [18]. As they are designed for bulk ATAC-seq data, running them on single-cell datasets is highly memory intensive. Therefore, for testing these methods we subset the hematopoietic cells dataset to only include 5 cell types (monocytes, B cells, CD8+ memory T cells, CD8+ naive T cells, natural killer cells) according to original annotations.

### 6.3 Simulation

Our simulation relies on varying the parameters from the hierarchical model in Section 3.3. There are 4 parameters needed to simulate data: 1) observation probability p_i_, 2) proportion of open cells *π*_j_, 3) background rate 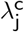, and 4) signal-to-noise ratio s_j_. In our simulations, we estimated p_i_ from the hematopoietic cells dataset to represent the sequencing depth variation between cells in real data. 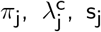 were either varied as hyperparameters for simulations shown in Fig. 4 or estimated from the hematopoietic cells dataset for simulations shown in Fig. 5. Below we show how parameters were estimated from data.

#### 6.3.1 Varying parameters in silico

For the simulations shown in Fig. 4, we only estimated p_i_ from the hematopoietic cells dataset (Section 6.3.2), while varying 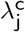 and s_j_ *in silico*. We fixed *π*_j_ = 0.3 for demonstration purposes, but our conclusions hold for other values as well (Fig. S2). To cover a dynamic range of parameter values, we simulated data with 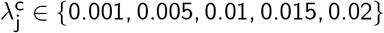 and s_j_ ∈ {2, 5, 10, 50, 100}. For each combination of 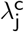 and s_j_, the simulation was repeated for 30 times and the mean AUROC is reported (Section 6.3.6). Similarly, for simulations shown in Fig. 5, each simulation was repeated for 30 times and the AUROC is reported.

#### 6.3.2 Estimating observation probability p_i_

Observation probability p_i_ was estimated from the hematopoietic cells dataset using the PIC model [4]. Below we adapt notations from Miao and Kim [4] to stay consistent with our previous definitions. Briefly, the PIC model introduces a binary vector T_i_ that indicates whether a genomic region j is measured in a cell i. Whether a region is measured depends on the observation probability q_i_ (Eq. 8),

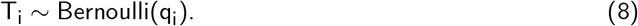

Although conceptually similar to our binomial measurement model (Eq. 5), the PIC measurement model assumes an ‘all-or-nothing’ mechanism—Tn5 insertion events are either all observed or all dropped out. Realistically, the more underlying insertion events there are in a region, the less likely all events in that region are dropped out. However, inference for p_i_ in Eq. 5 has no closed form solution and for data with generally low counts, q_i_ should be a good approximation for p_i_. Therefore, we used the get r by ct mat pq() function from the ‘PICsnATAC’ R package and used the estimated q_i_ as our observation probability p_i_. For the simulations shown in Fig. 4, we randomly sampled 10,000 observations from the estimated p_i_ to simulate from. For the simulations shown in Fig. 5, we only used the estimated p_i_ from relevant cell types (Section 6.3.4).

#### 6.3.3 Estimating background rate 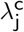

Background regions were used to infer 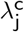. We chose background regions by using regions 500bp upstream and downstream of called peaks. Let k ∈ {1, …, M} index background regions. We assumed the same data generative process as our main model but all cells are closed in these regions, i.e. *π*_k_ = 0, such that all counts are due to background rate 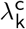. Then 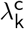 can be solved by matching the first moment. We denote 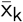 as the empirical mean of background region k and 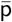 as the empirical mean of the previously estimated p_i_:

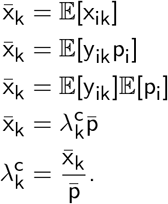

To model the effect of GC-content on background rate, we fit the following generalised additive model (GAM) using GC-content of each background region (GC_k_) as the predictor variable using ‘mgcv’ R package:

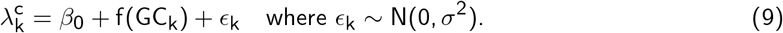

To prevent the fit from being affected by background regions with extremely high counts, background regions with 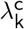 larger than 10 times the interquartile range were not used to fit the GAM. A total of 67,433 background regions (5.9% of all background regions) were filtered because of this reason. Lastly, to obtain an estimate for 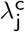 we use the fitted GAM to predict 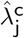 using the GC-content of called peaks (GC_j_).

#### 6.3.4 Estimating proportion of open cells *π*_j_

We subset data from 3 dominant cell types in the hematopoietic cells dataset according to annotations by the original authors, which include T cells (both naive and memory), B cells, and monocytes, resulting in a total of 7,884 cells in this subset of data. As we focus on simulating marker peaks for these cell types (Section 6.3.5), we assumed these peaks are only ‘open’ in their respective cell types, therefore we estimated *π*_j_ to be their cell type proportions, respectively 0.385, 0.378, and 0.237.

#### 6.3.5 Estimating signal-to-noise ratio s_j_

To simulate peaks with relevance to biology, we estimated parameters from peaks that are strongly associated with a curated set of marker genes (Table 4). We chose marker genes specific to the 3 chosen cell types based on literature [36, 37]. For each marker gene, there are multiple peaks within 500bp of the gene body. Among these peaks, we selected the peak that has the highest mean count to estimate our parameters from (Table 4).

Estimation of the signal-to-noise ratio s_j_, since all the other parameters are already estimated, can be achieved by matching the first moment. Again, we denote 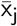 as the empirical mean of peak j and 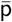 as the empirical mean of the estimated p_i_:

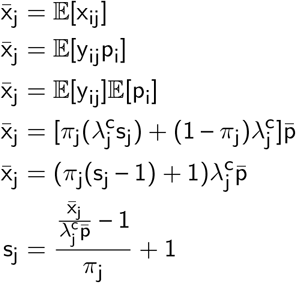

#### 6.3.6 Computation of posterior probability

The posterior probability of cell i being ‘open’ in region j is given by:

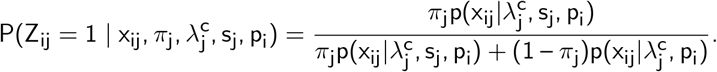

The marginal p.d.f of x_ij_ is given by:

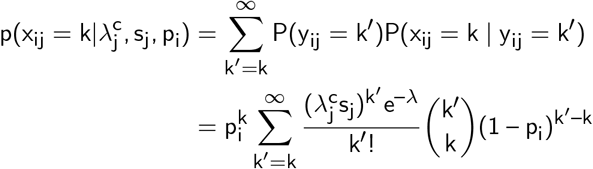

In a realistic scenario, the parameters should be estimated from data to calculate the posterior. However, in our simulations, we calculated the posterior with ground truth parameters (i.e. assuming perfect recovery of parameters). Note that the integral for the marginal p.d.f has no closed form solution. Therefore, for computational purposes it is approximated by addition up to 50. Further note that for the marginal p.d.f of the closed component, s_j_ = 1.

#### 6.3.7 Computation of AUROC

The AUROC is calculated using the ‘pROC’ (v1.18.5) [38] package in R with default parameters. We used simulated chromatin states of each cell as response and the posterior probability as the predictor.

### 6.4 Clustering analysis

The clustering analysis (Fig. 5) is performed using the Signac (v1.13.0) [6] pipeline with default parameters. Briefly, for each selected quantile of peaks, TF-IDF normalisation followed by Singular Value Decomposition (SVD) is performed to reduce the dimensions of the data. Then, UMAP is calculated with the first 2 to 30 LSI components, dropping the first component as recommended. Neighbour purity is calculated with the neighborPurity() function in ‘bluster’ R package (v1.16.0) [25], with the first 2 to 30 LSI components as input.

## 7 Data and code availability

Data and code to reproduce the figures in this manuscript are available at the following Github repository: https://gitlab.svi.edu.au/biocellgen-public/gath_2023_scatac_mixture_modelling_reproducibility.

## 9 Acknowledgements

The authors acknowledge Jeffrey Pullin and Sagrika Chugh for helpful discussions about this manuscript. This work was supported by NIH R01HG011886 and NHMRC GNT1195595 to DJM.

