## Supplementary Figures for "Going beyond cell clustering and feature aggregation: Is there single cell level information in single-cell ATAC-seq data?"

### Supplementary information

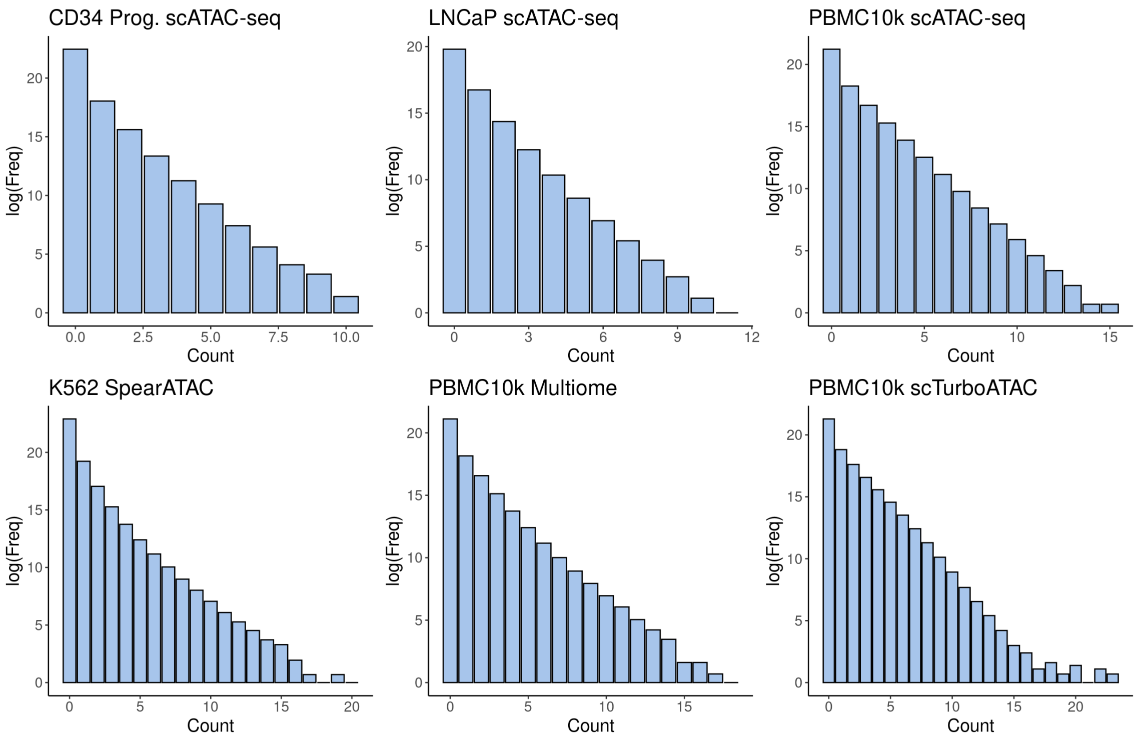


Figure S1: Histogram of paired-insertion counts (PIC) for all datasets analysed.

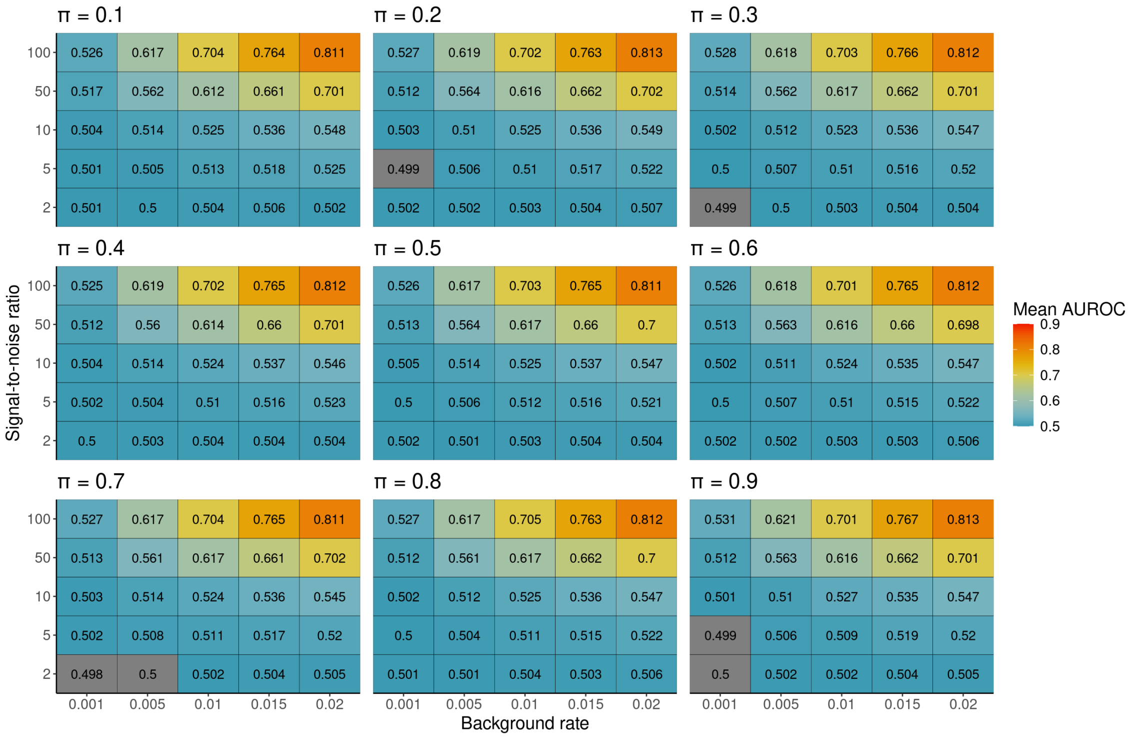


Figure S2: Simulations with varying $\pi$.

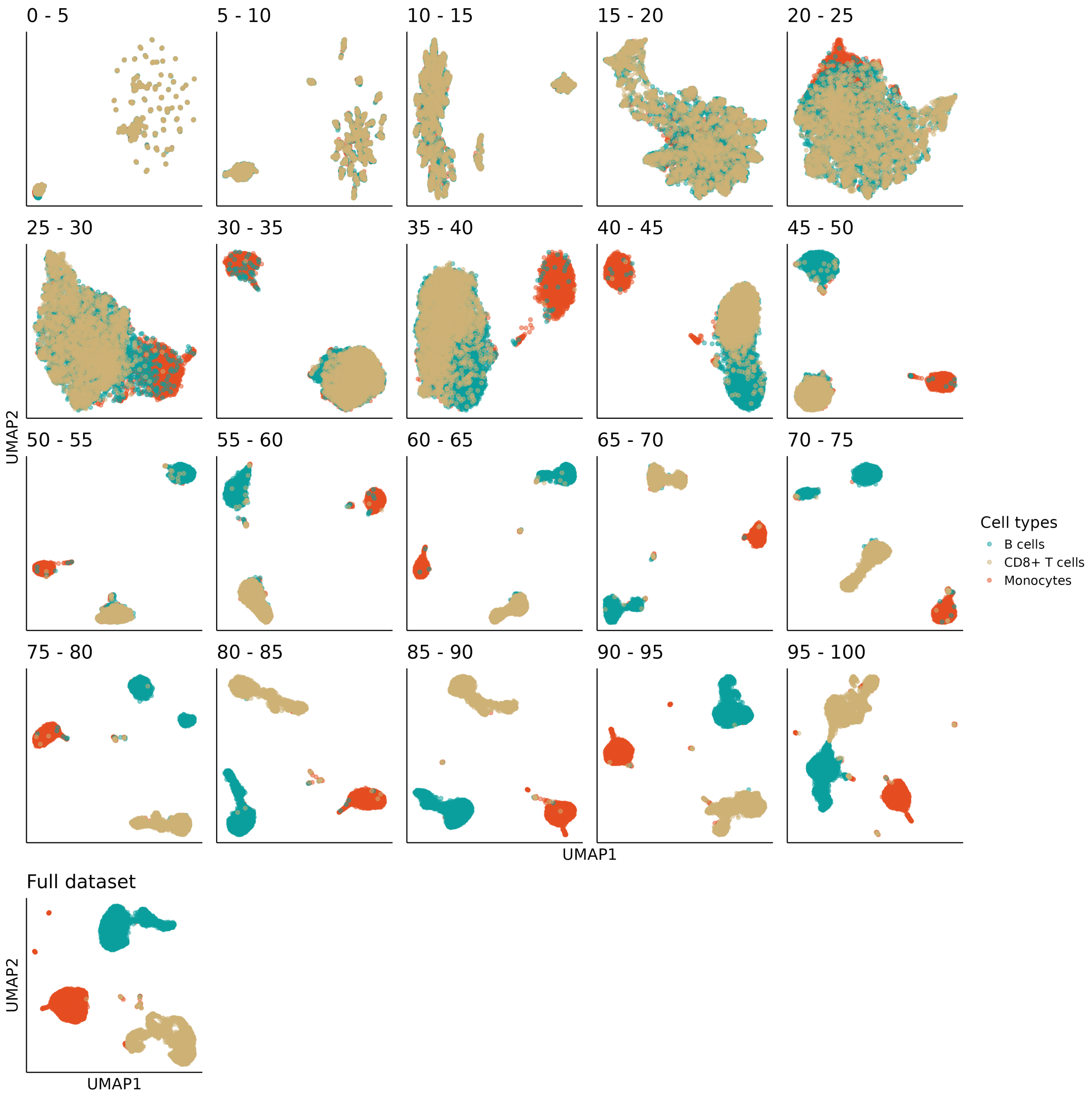


Figure S3: UMAPs of clustering with varying peak quantiles.
